# CentIER: accurate centromere identification for plant genomes with sequence specificity information

**DOI:** 10.1101/2023.12.24.573279

**Authors:** Dong Xu, Huaming Wen, Wenle Feng, Xiaohui Zhang, Xingqi Hui, Yun Xu, Fei Chen, Weihua Pan

## Abstract

Centromere identification is one of the important problems in genomics, providing a foundation for the studies of centromeres in aspects of composition, functionality, evolution, inheritance, and variation. The existing wet-experiment-based method is costly and time-consuming, while the bioinformatic method can only detect tandem repeats losing non-repetitive sequence regions in the centromere. To address these shortcomings, we introduce a new pipeline, CentIER, for the automatic and accurate identification and annotation of centromere regions by taking advantage of the sequence specificity information. CentIER only requires users to input the genomic sequence, and then it can partition the centromeric region from a chromosome, identify tandem repeat monomers, annotate retrotransposons, and ultimately output visualized results. By referencing the experimentally determined centromere regions, it was discovered that the predictive accuracy of centromere recognition by CentIER exceeded 90%. Following the evaluation of CentIER’s accuracy, it was applied to investigate the sequence and distribution characteristics of centromeric retrotransposons and tandem repeat sequences of different species, providing insights into these traits in monocotyledonous and dicotyledonous plants.

## Introduction

Centromeres play an essential role in genome stability, cell division, and disease development. However, for a long time, due to the incomplete assembly of the genome and the high complexity of centromeric composition, complete and reliable centromeric sequences of eukaryotic genomes have not be obtainable, hindering deeper studies. Centromeric regions are characterized by the presence of a high proportion of extra-long tandem repeats (on the scale of Mb) with highly similar repeat units such as monomers and higher order repeats (HORs), which exceed the capability of genome assembly technologies based on short sequencing reads and normal long noisy reads (Melters et al., 2013). In recent years, PacBio HiFi and ONT ultra-long sequencing technologies and their corresponding genome assembly algorithms have significantly improved the completeness and accuracy of centromeric regions in a large number of eukaryotic reference genomes, providing an opportunity for deeper investigation of centromeres in aspects of composition, functionality, evolution, inheritance, and variation.

While all kinds of studies and analyses of centromeres need to start from the accurate identification of their sequences in high-quality (e.g., telomere-to-telomere (T2T), gap-free, gapless) genome assemblies, there is very little methodological research focusing on this problem. Only two existing methods currently exist for centromere identification. The wet-experiment-based method utilizes centromere-specific proteins (CenH3 in plants) to perform Chromatin Immunoprecipitation Sequencing (ChIP-seq) and maps centromeric reads to the assembly for identification, which is costly and time-consuming (Liu et al., 2023; Marques et al., 2015). The bioinformatics method detects all tandem repeat monomers (TRs) in the whole genome first, and performs the identification by assuming the centromeric TRs are the most abundant among all TRs of a genome (Shi et al., 2023). For example, quarTeT can be used for identifying centromeric repeats (Lin et al., 2023). Although this assumption is valid in most situations, this method has two serious shortcomings as follows. Firstly, the bioinformatics method can only detect TRs in centromeres, while the entire centromeric regions also contain interspersed repeats such as long terminal repeat retrotransposons (LTR-REs) and non-repetitive genomic regions (Liu et al., 2015). Secondly, non-centromeric regions may also contain highly repetitive sequence units, leading to a certain amount of false-positive identification.

In this study, we initially conducted a comprehensive analysis to elucidate the shortcomings in current methods for centromere identification. Subsequently, an innovative algorithm was proposed, which integrates information on *k*-mers (substrings of length k) and TRs to accurately identify centromere regions. This groundbreaking approach resulted in the development of a novel pipeline known as CentIER. To verify its accuracy, CentIER was rigorously validated using different plant genomes (such as *Arabidopsis thaliana* and *Oryza sativa*). Additionally, the application of CentIER was extended to the recognition of centromeres in diverse species, enabling a characterization of the compositional features of plant centromeres. To the best of our knowledge, CentIER currently stands as the first program capable of identifying complete centromeres, distinguishing itself from quarTeT used for identifying centromeric repeat regions.

## Results

### Some difficulties exist in the current methods for identifying the centromere

The conservation of the CenH3 protein plays a pivotal role in determining the applicability of experimental methods for identifying centromeric regions in plants (Chen et al., 2023). The conservation of CenH3 proteins across various plant species was investigated by downloading protein sequences from NCBI and constructing a phylogenetic tree. Phylogenetic analysis demonstrated species-specific differentiation of CenH3, with a trend for CenH3 proteins from closely related species to cluster together on the same branch (Figure S1, the red dashed boxes). Sequence diversity analysis was performed on two CenH3 proteins (derived from *O. sativa* and *Brassica rapa*, respectively), which exhibited the greatest degree of differentiation on the phylogenetic tree (Figure S1). The result demonstrated that the sequence similarity between OsCenH3 (from *O. sativa*) and BrCenH3 (from *B. rapa*) is merely 60.2%, with the N-terminal sequence exhibiting the lowest degree of similarity (Figure 1A). In other words, due to the variations in CenH3 proteins across different species, employing the same CenH3 protein antibody in ChIP-seq experiments may potentially impede the accurate identification of the centromere region, thereby increasing the challenge of identifying the centromere region using experimental methods. Therefore, it is crucial to develop accurate bioinformatics prediction methods for the identification of centromeres in a broader range of plant species.

**Figure 1.**
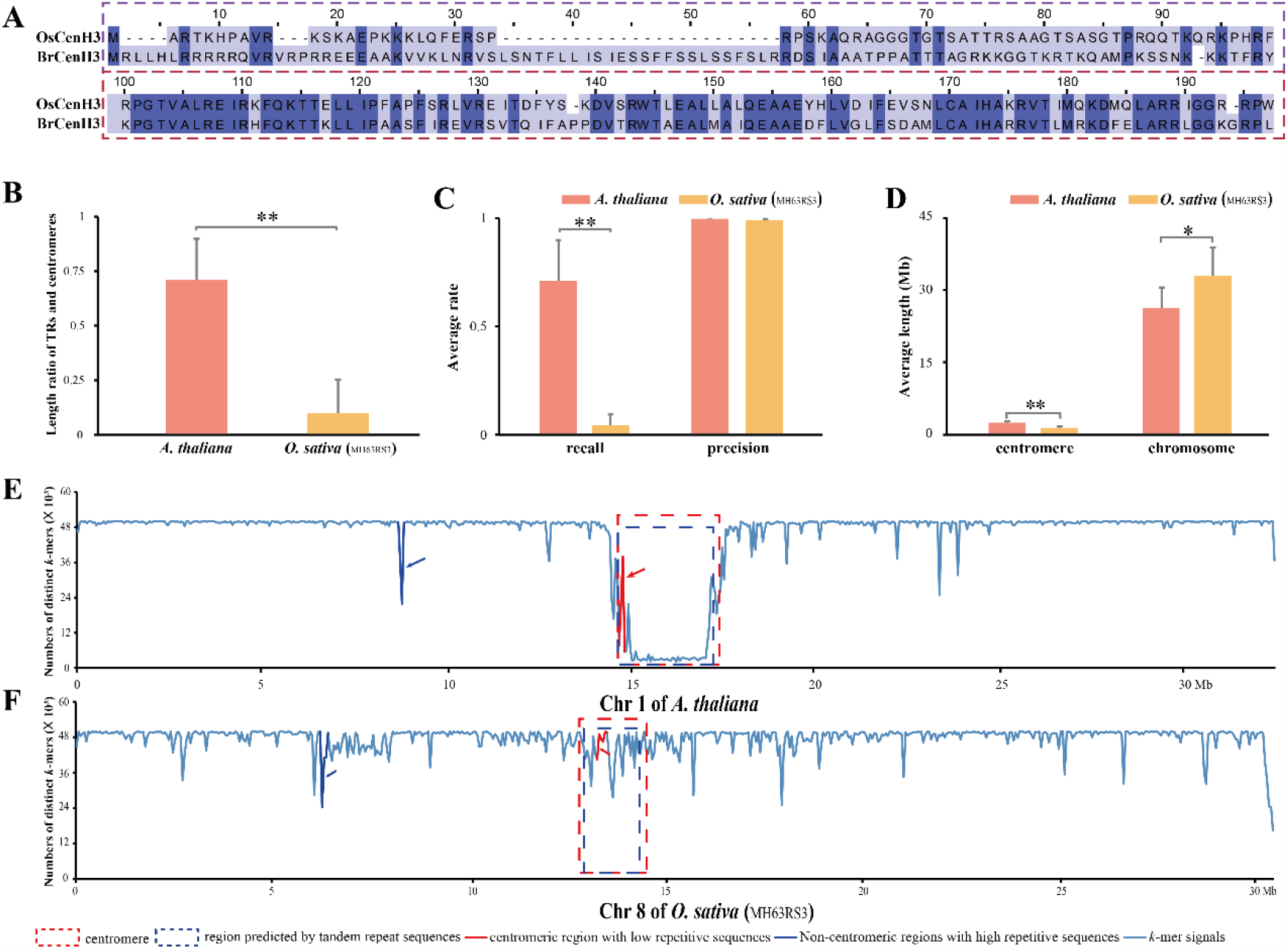
Sequence characteristics associated with centromeres. (**A**) Sequence alignment between OsCenH3 and BrCenH3. (**B**) The ratio between the length of intervals containing TRs and the length of centromeres. (**C**) The recall rate and precision rate between the regions predicted by TRs and centromeric regions. (**D**) The average lengths of chromosomes and their corresponding centromeres from *A. thaliana* and *O. sativa*. The *k*-mer signal distributions of the *A. thaliana* chromosome 1 and the *O. sativa* chromosome 8 were respectively depicted in (**E**) and (**F**). The red dashed boxes, centromeres; The red arrows, the regions in centromeres with normal *k*-mer signal level; The blue arrows, the non-centromere regions with abnormal *k*-mer signal level. Mb, Megabase; chr, chromosome.

The utilization of Tandem Repeats Finder (TRF) for querying TRs is a widely employed method in bioinformatics for identifying centromere regions (Deng et al., 2022; Shi et al., 2023). The accuracy of this particular approach is contingent upon the size and chromosomal positioning of the intervals encompassing tandem repeats and centromere regions. The ratio of the lengths between the centromere regions, known in *A. thaliana* and *O. sativa* (Naish et al., 2021; Song et al., 2021), and their respective TRs regions was examined in this study. It was observed that the lengths of the TRs regions in both species were smaller than those of the centromere regions, and there was a significant difference in the length ratio. (Figure 1B, *p*-value <= 0.01). Subsequently, the recall ratio and precision ratio of the TRF method were calculated separately (see **Methods** section for calculation details). It was observed that the recall ratio of both species was found to be less than 1 with significant interspecies differences (*p*-value <= 0.01). This suggests that the TRs regions tend to be smaller than the centromere regions. Conversely, the precision ratio approached 1 (Figure 1C) indicating that TRs regions can effectively locate centromere regions. Meanwhile, the chromosome lengths and centromere lengths were investigated in both species. The results revealed significant differences in centromere lengths and chromosome lengths among different species (Figure 1D, *p*-value <=0.05). It is worth noting that the elongation of chromosomes does not inherently imply an extension in the length of centromeres.

### Centromeric regions exhibit significant differences from other chromosomal regions in the profile of sequence specificity

Based on the analysis above, it is necessary to incorporate additional methods to rectify and enhance the accuracy of the bioinformatics-based prediction of centromeres. Since centromeres are mainly composed of repeat sequences, it is reasonable to speculate that the sequence specificity of centromeric regions is lower than other chromosomal regions. The genomes of *A. thaliana* and *O. sativa* (MH63RS3 and ZS97RS3 cultivars) with known centromere regions were examined to substantiate our hypothesis. More specifically, the chromosomes were cut into small segments (50,000 bp), and the number of distinct *k*-mers (genomic substrings of length *k, k*=21 in the experiments) per segment was used as *k*-mer signal of sequence specificity.

The signals (numbers of distinct *k*-mers) in non-repetitive genomic regions per segment were over 49,000 (the number of all *k*-mers per segment, Figure 1E and F) which was conserved across different species showing an extremely-high sequence specificity and universality across species (Figure S1), while the ones in repetitive regions were significantly lower (Figure 1E and F, the blue arrows). Compared to other repetitive regions showing comparatively short low-signal regions in the profiles, centromeres are the extra-long recessed regions in chromosomes with continuous low-signals (Figure 1E, the red dashed box), and this phenomenon can be observed on most chromosomes (Figure S2). Meanwhile, it is possible to find other repetitive sequences flanking the tandem repeat regions in the centromeric regions (Figure 1E, the regions between the blue dashed box and the red dashed box). Furthermore, although the signals of some non-repetitive regions in the centromere may be higher than the tandem repetitive regions, their signals are still much lower than normal chromosomal regions (Figure 1E, the red arrow). Therefore, the centromere regions can be accurately determined by querying low *k*-mer signals. However, it has been observed that a minimal fraction of centromeres may exhibit signal levels similar to those of other genomic regions (Figure 1F and Figure S1). In an effort to more accurately predict centromere regions, new algorithms and programs are being developed that consider the combination of sequence specificity and TRs information.

### CentIER algorithm and tool

According to the above observations, we developed CentIER algorithm for centromere identification on genome sequence. CentIER algorithm is the main module of CentIER tool, and it contains three steps as follows (Figure 2B). First, CentIER detects the abnormal (low-signal) regions in the profile of sequence specificity (Figure 2D, the red lines). Since a fixed threshold is difficult to set for distinguishing abnormal regions containing complete centromeric sequences from the normal regions containing no centromeric sequences on different genomes, we designed an adaptive algorithm for deciding this threshold for each genome according to its whole-genome sequence-specificity information. Second, since centromeres contain non-repetitive sequences with comparatively-high-signal in the profile of sequence specificity, in many situations, they are broken into discontinuous abnormal regions and these non-repetitive sequences are misclassified into normal regions. To solve this problem, CentIER merges the detected abnormal regions if the length of the normal regions between them is within a tolerance threshold (several potential abnormal regions may thus be produced) (Figure 2D, the orange dotted boxes). This key threshold affects the accuracy of identification to a large extent. A too-large threshold leads to the misclassification of pericentromeric repeats into centromere, while a too-small value leads to the identification of incomplete centromere. Third, since the merged abnormal regions include not only centromeric sequences but other tandem repeats, CentIER performs the filtering according to the copy numbers of TRs. Because, in most situations, centromeres are typically the regions on chromosomes that contain the highest number of TRs (Melters et al., 2013).

**Figure 2.**
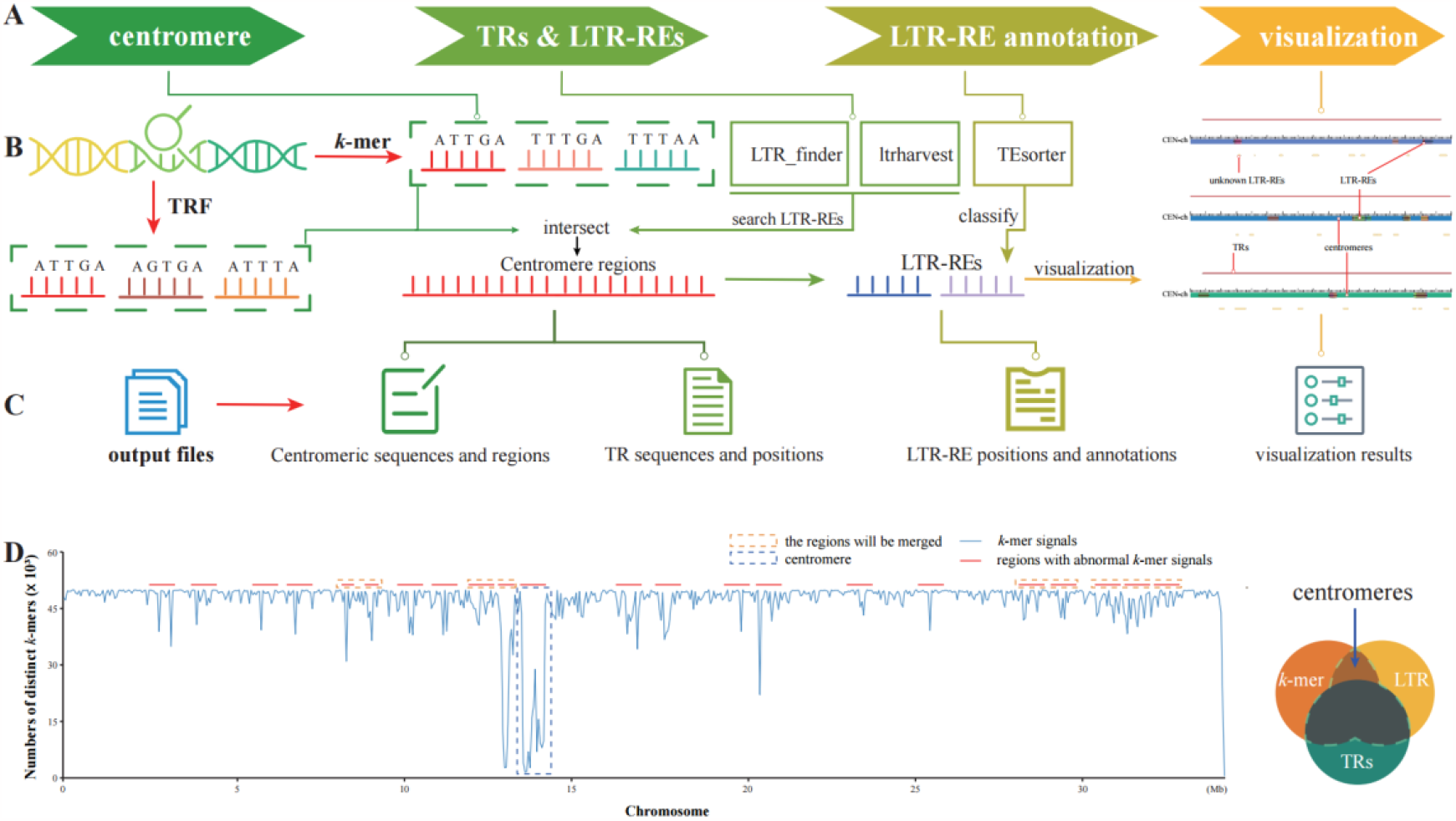
Overview of CentIER. Total four modules were set in CentIER (**A**). The workflow and the output results of CentIER were showed in (**B**) and (**C**), respectively. The *k*-mer signal of rice chromosome 11 was utilized as real data to demonstrate the method details about the identification of centromeres in CentIER (**D**).

In addition to the above utilization of repeat sequences, considering that the centromeric region is specifically distributed with some LTRs, we also counted the number of LTR distributions to select the region with the highest number of LTRs among multiple candidate regions as the centromeric region. In summary, CentIER recognizes the region with the highest number of TRs repeats, queries for *k*-mer abnormal regions that intersect with this TRs region, and expands the TRs region using these *k*-mer regions. There may be multiple candidate regions generated. Subsequently, we detected the number of LTRs in these candidate regions and selected the region with the highest number of LTRs as the centromeric region. In addition to CentIER algorithm, CentIER tool provides some auxiliary functions of downstream analyses such as the identification of TRs, identification and annotation of LTR-REs, and visualization (Figure 2A and C).

### Validation of CentIER accuracy

Centromere regions of *A. thaliana, O. sativa* (MH63RS3 and ZS97RS3) and *Z. mays* with T2T level genomes have been identified by experimental evidences (Chen et al., 2023; Naish et al., 2021; Song et al., 2021). Therefore, we selected them as the real datasets for testing the accuracy of CentIER. To the best of our knowledge, the quarTeT is currently the only program available for centromeric repeat identification in T2T genomes (Lin et al., 2023). Therefore, we intend to compare the prediction results of CentIER with those of quarTeT.

The results were showed in Figure 3. The genome of *A. thaliana* served as our initial testbed to evaluate the prediction accuracy of CentIER. We found that the majority of centromere sequences from all chromosomes were accurately predicted by CentIER (Figure 3A). The CentIER accuracy of centromere prediction in diverse species was assessed by utilizing the remaining three genomes and using the parameters of predictive accuracy, recall rate, precision rate as well as F1-score (definition and algorithm of the four parameters reference **Methods** section), and the results were compared against the predictions made by quarTeT (Figure 3 B-E). The predictions made by CentIER successfully identified all centromeres of the three species (two rice varieties and *Z. mays*), either in their entirety or partially, exhibiting a 21% improvement over quarTeT ’s predictions (Figure 3B). CentIER achieved an average recall rate of 63% and precision rate of 73% in its prediction results, surpassing quarTeT ’s predictions by 26% and 13%, respectively (Figure 3C and D). Meanwhile, the average F1-score of CentIER was also 18 % higher than that of quarTeT (Figure 3E). These results highlight the higher accuracy of CentIER over quarTeT and its potential for accurate centromere prediction in various species. Building on this premise, the centromere regions of different plants with T2T-level genomes were predicted and subjected to analysis using CentIER, resulting in the subsequent findings.

**Figure 3.**
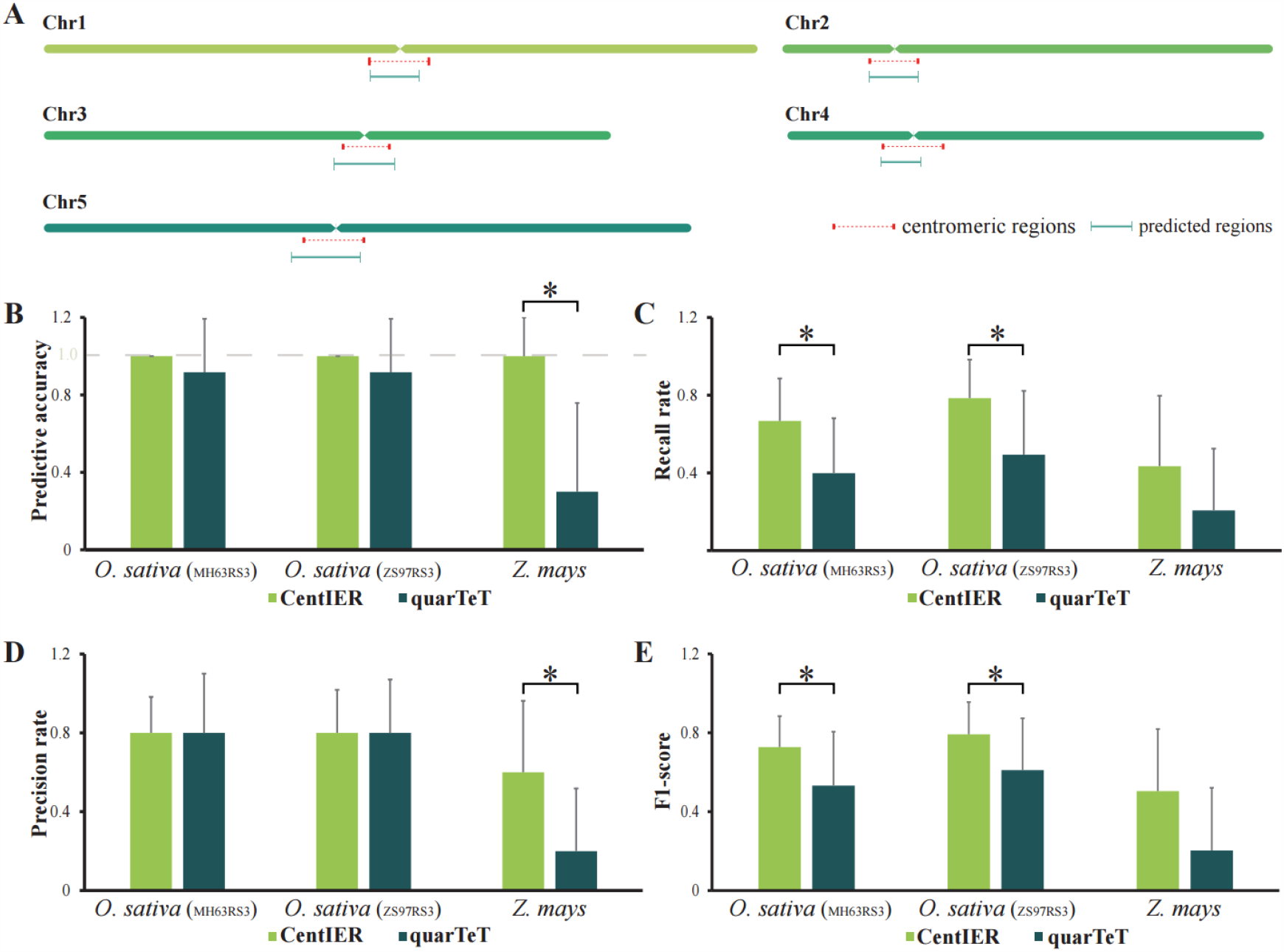
Accuracy of CentIER. CentIER effectively anticipated the majority of centromeres in *A. thaliana* (**A**). (**B-E)** illustrated the values for predictive accuracy, recall rate, precision rate, and F1-score. *, *p*-value <= 0.05; *O. sativa* (MH63RS3), the rice cultivar MH63RS3; *O. sativa* (ZS97RS3), the rice cultivar ZS97RS3.

### Centromeres affecting haplotype length but not the main reason

Changes in the number of TRs repeats and the random insertion of LTR-REs can result in changes to centromere length. Meanwhile, we observed that haplotypes of the same chromosome are typically different in length (Han et al., 2023). However, the factors leading to haplotype length variation are still unknown. As far as we know, the genome of *Actinidia chinensis* stands out as the sole T2T-level genome known to possess multiple haplotypes, with the exception of regions containing unknown sequences. We thus downloaded genomic sequences of two haplotypes of *A. chinensis* (Han et al., 2023) and used CentIER to analyze centromeres, to determine whether TRs and LTR-RTs could significantly affect haplotype length. The statistical results are shown in Figure 4.

**Figure 4.**
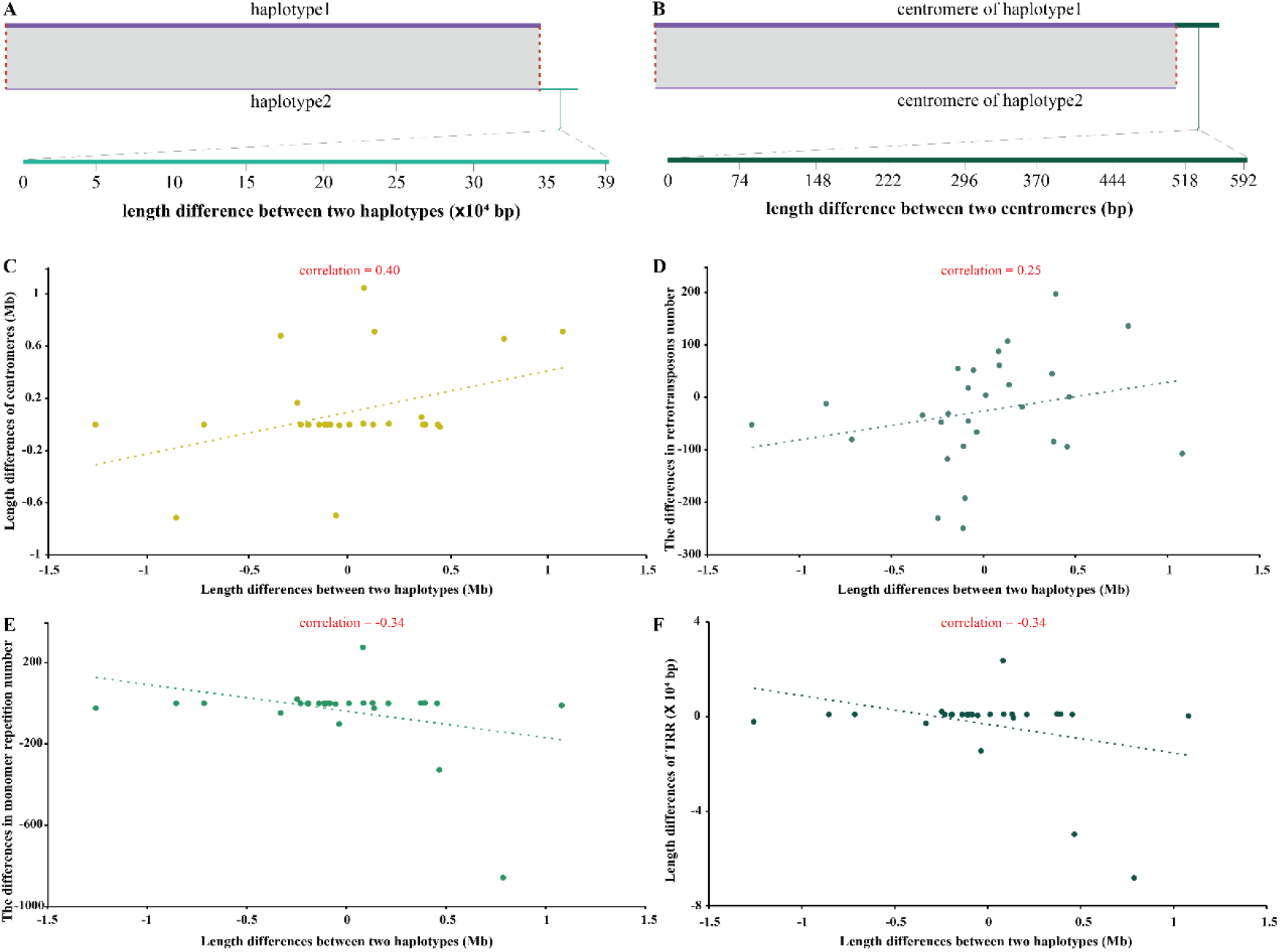
The relationship between centromeres and haplotypes. Taking Chromosome 18 of *A. chinensis* as an example, haplotype 2 is longer in length compared to haplotype 1 (**A**), but the centromere of haplotype2 is shorter than that of haplotype1 (**B**). Figures **C** through **F** illustrated the relationships between centromere length differences, the number of retrotransposons, the number of TRs, and the difference in TRR length, and the lengths of two haplotypes. ‘Length differences between two haplotypes’ refers to the difference in length obtained by subtracting haplotype 1 from haplotype 2.

Taking Chromosome 18 as an example, we observed that the length of haplotype 1 was shorter than that of haplotype 2 (Figure 4A). Conversely, the centromeric region of haplotype 1 was longer than that of haplotype 2 (Figure 4B). Put differently, an increase in centromeric region did not lead to a corresponding increase in haplotype length. To strengthen the conclusion, we evaluated all haplotypes and centromeres of *A. chinensis*. The analysis revealed a positive correlation between centromere length and haplotype length (Figure 4C). However, it is worth noting that the correlation coefficient between them was only 0.4, which is significantly less than 1. Our findings suggest that while there is a tendency for the length of the centromere to influence the length of the haplotype, it is not a strong or decisive factor in determining haplotype length. Building upon this, we further conducted additional research to explore the potential impact of the centromere’s composition on haplotype length variations. The correlation coefficients were 0.25 (for retrotransposon), -0.34 (for monomer repeat) and -0.34 (for the product of the repetition count and tandem repeat (TRR)), which were not sufficient to support a strong relationship between them and the length difference of the haplotype (Figure 4D-E). Hence, we concluded that while alterations in centromere length may contribute to variations in haplotype length, they were not the predominant factor responsible for the observed discrepancies in haplotype lengths.

### Sequence characteristics and distribution differences of TRs and LTR-REs in monocotyledonous and dicotyledonous plants

Although the centromeric information of several species were reported in previous studies, the sequence characteristics and distribution differences of TRs and LTR-REs need to be further analyzed and summarized between monocotyledonous and dicotyledonous plants. A total of 58 LTR-REs belonging to two categories were detected in four dicots (*A. chinensis, A. thaliana, Citrus limon* and *Vitis vinifera*) and two monocots (*Erianthus rufpilus* and *O. sativa*) (Shi et al., 2023; Wang et al., 2023). The abundance of LTR-REs in dicots was higher than the abundance of LTR-Res in monocots, but the difference did not reach a significant level (*p*-value >= 0.05, Figure 5A). Only Tat I, Tat II and Tat III in 58 LTR-REs were not detected in the two monocots while the quantity of Bel-Pao, Helitron and Maverick in dicots is significantly higher than these in monocots (Figure 5 A and B, *p*-value <=0.05). Over 82 % of class I LTR-REs and 75 % of class II LTR-REs can be detected in this experiment. In summary, despite the detection of a large number of LTR-REs in dicots or monocots, the distribution of LTR-REs does not seem to be significantly different between dicots and monocots.

**Figure 5.**
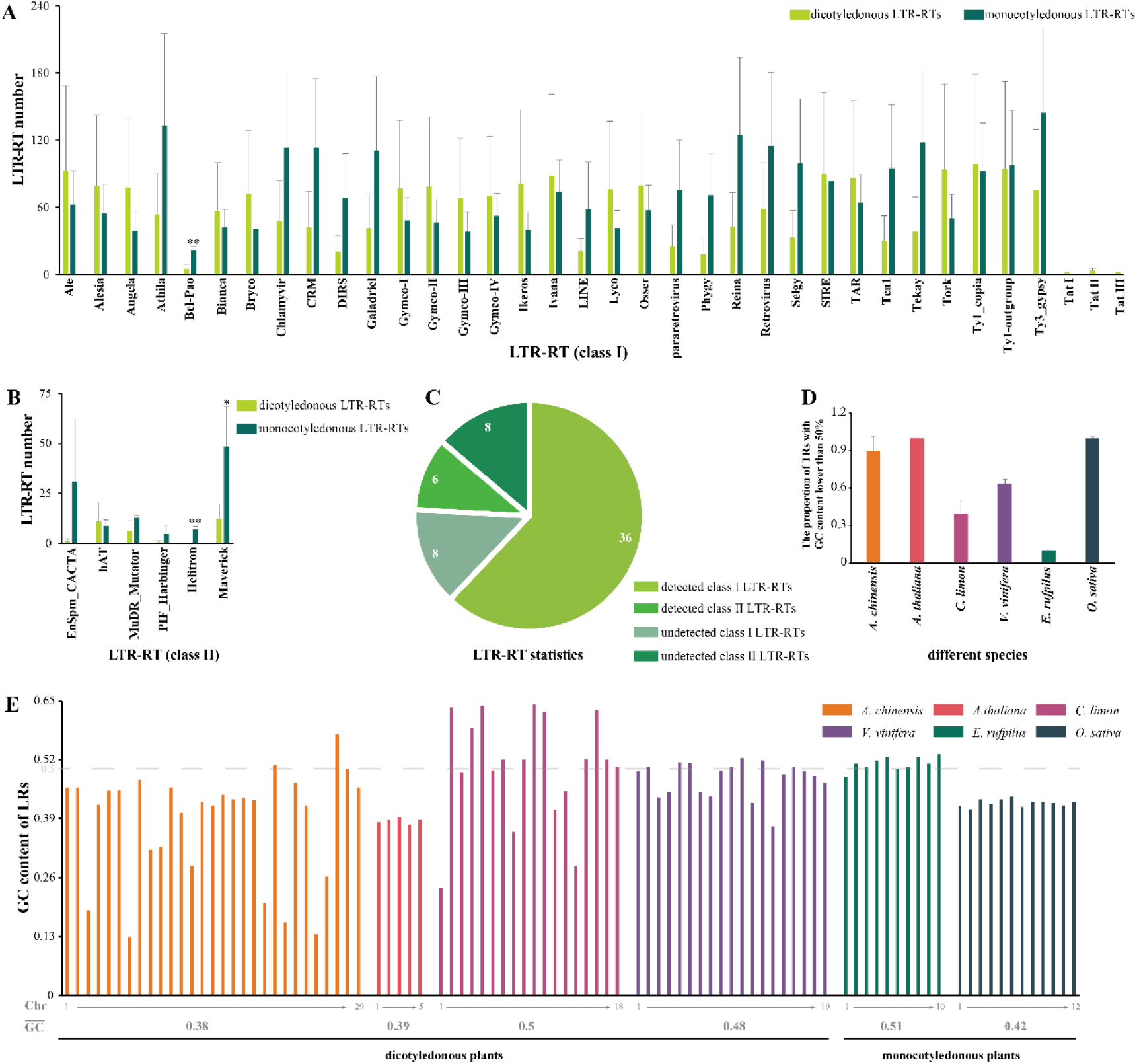
Sequences and distribution characteristics of TRs and LTR-REs in monocotyledonous and dicotyledonous plants. Distribution of LTR-RE quantities in monocotyledonous and dicotyledonous plants (**A** and **B**). (**C**) LTR-RE statistics. **(D)** The proportion of TRs with GC content lower than 50 % among in monocotyledonous and dicotyledonous plants. **(E)** GC content of chromosomal LTR-REs in monocotyledonous and dicotyledonous plants. Chr, chromosome. 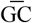, the average of GC content.

The finding that TRs have lower GC (G, Guanine; C, Cytosine) content has been reported in several animal studies (Talbert and Henikoff, 2020). The validity of this conclusion in plants and whether there are differences in GC content between monocotyledonous and dicotyledonous plants remain to be determined. We calculated the GC content of TRs on each chromosome of the six species mentioned above (Figure 5E), and conducted statistics on TRs with a GC content below 50% as shown in Figure 5D. We found that the average GC content of TRs in *C. limon* and *E. rufpilus* was equal to or greater than 0.5 (Figure 5E). The average GC content of TRs in *A. chinensis* was the lowest (at 0.38) among these six species, while the GC content of TRs in other species was below 0.5 (Figure 5E). The length and sequences of TRs may vary among different chromosomes leading to the differences in GC content. For example, most of the *A. chinensis* chromosomes have TRs with GC content below 0.5. However, the GC content of TRs on Chromosome 27 was higher than 0.5. We calculated the proportion of TRs with GC content below 50% in each species and found that the proportion is highest in *A. chinensis* (dicot), *A. thaliana* (dicot) and *O. sativa* (monocot), which approached or exceeded 90% while the proportion in *C. limon* (dicot) and *E. rufpilus* (monocot) was lower than 50% (Figure 5D). In a nutshell, the GC content of TRs in plants may be higher than 0.5, while there were no significant differences in the GC content of TRs between monocotyledonous and dicotyledonous plants.

## Discussion

### No significant differences of centromeric components between monocots and dicots

Our experiments provide further evidences that there are no significant differences in centromeric composition between monocotyledonous and dicotyledonous plants. Firstly, we observed no significant differences in the types of LTR-REs. Secondly, with the exception of a few LTR-REs, the majority of LTR-REs do not exhibit significant differences in quantity between monocotyledonous and dicotyledonous plants (Figure 5A and B). Consistent with previous studies (Barbosa et al., 2021), our results showed a lack of GC content in centromeric sequences in most eukaryotes (Figure 5E, *A. thaliana* and *O. sativa*). Additionally, our study demonstrated that the average GC content of TRs in *C. limon* and *E. rufpilus* was equal to or greater than 0.5, indicating that GC content exceeding 50% could occur in both monocotyledonous and dicotyledonous plants. Our findings provided additional evidence supporting the notion that centromeres seemed to exhibit a high degree of randomness in their sequence composition. Further investigation is warranted to unravel the underlying reasons or driving forces behind the observed random variation in centromere composition.

### Potential factors affecting the accuracy of CentIER

Based on our experimental results, CentIER demonstrates a high level of accuracy in centromere prediction. However, it is important to acknowledge that the prediction accuracy of CentIER can be influenced by unique genomic sequence features. The abundance of TRs within the genome serves as a critical factor that significantly impacts the prediction accuracy of CentIER. In one scenario, TRs were not distributed in the centromeric region. For example, the absence of TRs was observed in chromosomes 5 and 8 of maize (Chen et al., 2023). In another scenario, TRs were present in the centromeric region, but the number of TRs was too low to distinguish the centromeric region from other repetitive regions in the chromosomes (such as telomeres). For instance, no significant decrease in *k*-mer signal was observed in the regions of chromosomes 10 of watermelon (Figure S3)(Deng et al., 2022). As shown in Figure 5E, the distribution of LTR-Res on chromosomes exhibits a high degree of randomness. If TRs are not informative, it is truly hard to find other DNA sequence features to accurately identify the centromere regions. However, this may not be the primary reason affecting the accuracy of CentIER, because for the majority of eukaryotic genomes, their chromosomes should contain a substantial number of TRs (Melters et al., 2013; Presting, 2018). If the majority of chromosomes in a T2T-level genome lack or possess only a limited number of centromeric TRs, it suggests a probable occurrence of assembly-related challenges during centromere assembly. From this standpoint, CentIER could be considered a potential approach for assessing the quality of genome assembly.

As mentioned in the **Introduction**, the assembly of complete centromeres poses a significant challenge in genome assembly due to the presence of highly repetitive sequences. The prediction accuracy of CentIER may be compromised if centromeres are either not fully assembled or only partially assembled due to limitations in assembly or sequencing techniques. In essence, factors that affect the integrity of centromere assembly have the potential to impact the accuracy of CentIER in centromere identification. Additionally, it is important to note that CentIER was primarily designed based on the sequence characteristics of plant centromeres, and its effectiveness in predicting centromeres in animals and other species requires further validation.

### In the future

The utilization of TRs to identify centromere regions is currently the primary basis for centromeric identification using bioinformatics approaches (Fu et al., 2023; Zhou et al., 2023). According to our results and previous studies, we found that some significant variations in the sequence of TRs across different chromosomes can be detected even within a same genome (Melters et al., 2013). Furthermore, there is no observed difference in the distribution of LTR-REs among different species (Figure 5A). In other words, these evidences suggests that there seem to be few DNA sequence features that can be used for centromere identification. Indeed, we attempted to learn the characteristics of centromere regions through deep learning using LSTM models. However, due to limited data availability (limited number of T2T-level genomes currently accessible) or other factors, we did not achieve a satisfactory centromere recognition model. In upcoming research, we aim to explore other features that may be associated with centromere specificity to further enhance prediction accuracy.

## Methods

### Phylogenetic analysis and sequence alignment

A total of 47 CenH3-like proteins from different plant species were downloaded from NCBI (https://www.ncbi.nlm.nih.gov/gene). It was discovered that multiple CenH3-like proteins from the same species may have been included in the NCBI database, and their sequences exhibited a high degree of similarity. Therefore, only the CenH-like protein with the longest length from the same species was selected for the construction of the phylogenetic tree. The phylogenetic tree was built by using MEGA software with 1,000 bootstrap computations. The sequence alignment between *OsCenH3* and *BrCenH3* was performed by the Mafft function of SPDE (Xu et al., 2021) and using Jalview for alignment visualization (Waterhouse et al., 2009).

### Dataset download and usage of tools

The genomic data with complete sequences used in this study was downloaded from the following websites, respectively:

*A. thaliana* (Naish et al., 2021), https://github.com/schatzlab/Col-CEN/tree/main/v1.2;

*A. chinensis* (Han et al., 2023), http://182.92.183.62/;

*C. limon* (Bao et al., 2023), https://ngdc.cncb.ac.cn/search/?dbId=gwh&q=GWHCBFQ00000000.1;

*E. rufpilus* (Wang et al., 2023), http://sugarcane.zhangjisenlab.cn/SugarcaneDB/#/downloads;

*O. sativa* (Song et al., 2021), http://rice.hzau.edu.cn/cgi-bin/rice_rs3/download_ext;

*V. vinifera* (Shi et al., 2023), https://zenodo.org/record/7751391#.ZBgVmcJBy3A;

*Z. mays* (Chen et al., 2023), https://www.ncbi.nlm.nih.gov/bioproject/?term=PRJNA751841.

All parameters were set to their default values when using the tools (including CentIER and quarTeT) for centromere identification and analysis in the mentioned species.

### Adaptive algorithm for distinguishing abnormal regions in the profile of sequence specificity

The sequence is segmented into intervals of 10000 base pairs (bp) each, and the count of non-abundant *k*-mers within each interval is determined. Subsequently, these intervals are binned based on the number of non-redundant *k*-mers they contain. To distinguish normal intervals and abnormal intervals, we establish a threshold between normal bins and abnormal bins. This threshold ensures that the cumulative length of intervals in normal bins accounts for at least 85% of the total sequence length.

Considering the nature of centromeric regions, which may include non-repetitive segments amid repeat regions, we incorporate a tolerance level based on empirical observations. This tolerance factor allows for a potential number of normal intervals that may appear amid abnormal intervals to ensure the completeness of the centromere. To be specific, the final abnormal regions consist of consecutive abnormal intervals interspersed with normal intervals, within which the number of consecutive normal intervals must not exceed the defined tolerance level.

### Functions of downstream analyses in CentIER

In addition to CentIER algorithm, CentIER tool provides some auxiliary functions of downstream analyses such as the identification of tandem repeat monomers, identification and annotation of LTR-REs, and visualization. More specifically, LTR-finder (Xu and Wang, 2007) and ltrharvest (Ellinghaus et al., 2008) are utilized for confirming the positions of LTR-REs, and TEsorter (Zhang et al., 2022) is used to annotate the identified LTR-REs. Furthermore, all the important information including the centromeric regions, the TR and LTR-REs locations as well as the LTR-REs types can be visualized and outputted (Figure 2). All these operations can be completed in a fully automatic manner.

### Accuracy assessment

The parameters of predictive accuracy, recall rate, precision rate as well as F1-score were set in this study to assess the accuracy of CentIER. In detail, to evaluate the predictive accuracy, a scoring system was employed, wherein a score of 1 was attributed when the tool was able to predict any portion of the centromere sequence, while a score of 0 was given for unsuccessful predictions. The predictive accuracy was then computed by dividing the cumulative score by the total number of chromosomes.

We set A = the size of the centromeric region in the entire predicted region, B = the size of the whole centromere region, C = the size of the entire predicted region. The recall rate, expressed as the ratio between A and B, offers insights into the number of centromere regions that have been successfully predicted. The precision rate, represented by the ratio between A and C, indicates the proportion of predicted centromere regions out of the total predicted intervals. Interestingly, the recall rate and precision rate seem to exhibit an inverse relationship. Typically, as the recall rate increases, the precision rate tends to decrease. Consequently, to achieve a more comprehensive evaluation of these metrics, we utilized the F1-score, which is calculated using the following formula.

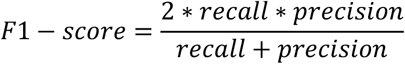

## Author contributions

W.P., F.C., Y.X. and D.X. conceived the study and designed the experiments. D.X. and H.W. developed the CentIER pipeline. D.X, W.F., X.Z. and X.H. collected the related data and performed the data analysis. W.P. and D.X. wrote the manuscript, while C.F. and Y.X. revised the manuscript.

## Data availability

The CentIER pipeline can be downloaded from https://github.com/simon19891216/CentIER/releases/tag/CentIERv1.1.

---

**Figure S1.**
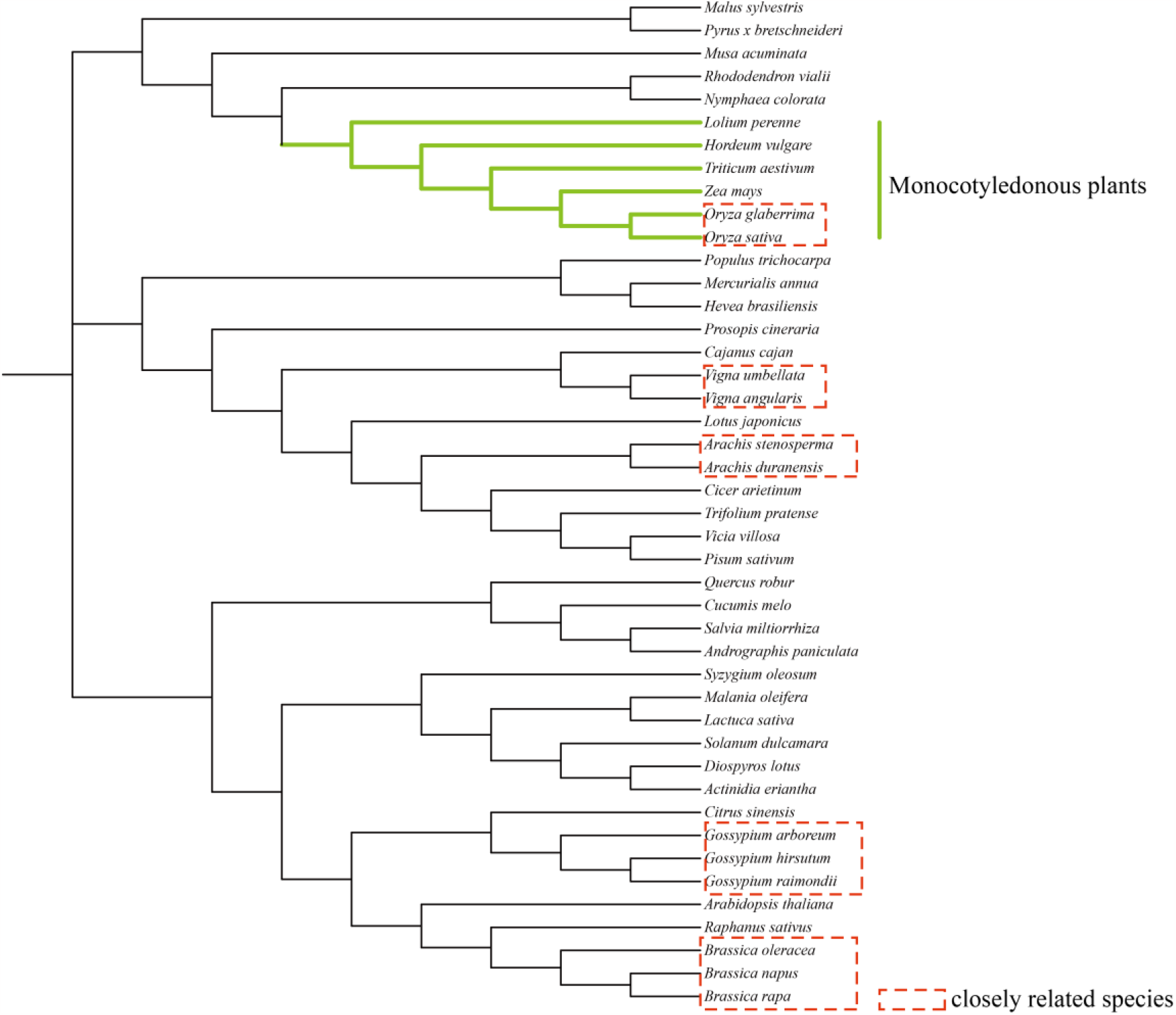
Phylogenetic tree constructed with CenH3 protein.

**Figure S2.**
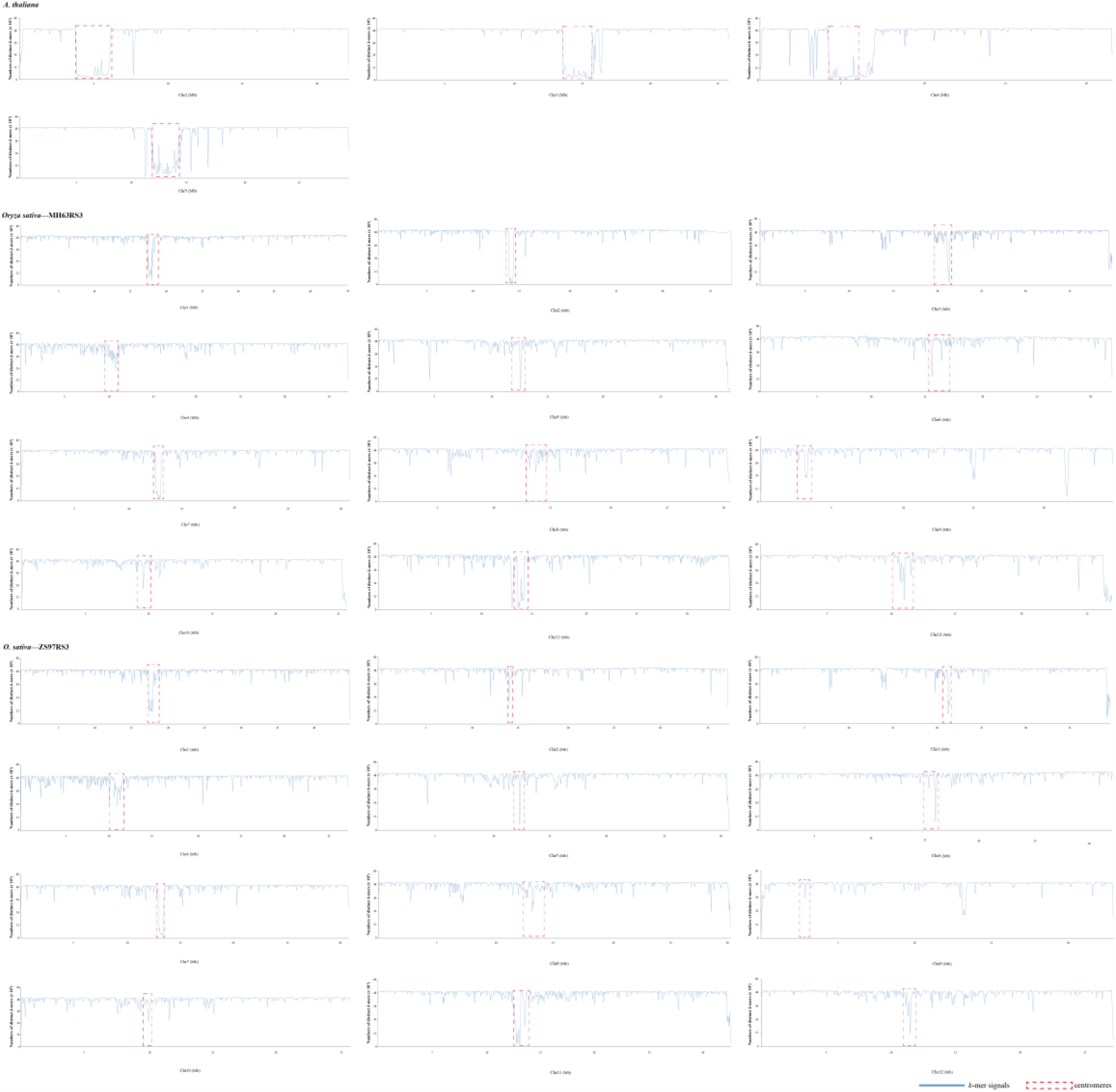
The *k*-mer distribution.

**Figure S3.**
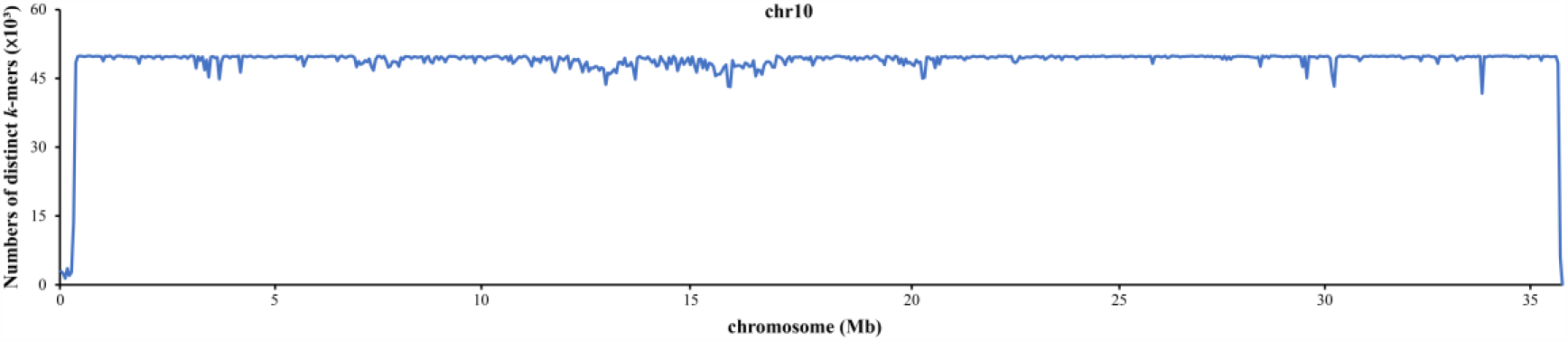
The *k*-mer distribution of Chromosome 10 of watermelon.

